# Heat stress drives transcription of LTR retrotransposons in the regenerative flatworm *Macrostomum lignano*

**DOI:** 10.1101/2024.12.18.629116

**Authors:** Kirill Ustyantsev, Stijn Mouton, Mikhail Biryukov, Jakub Wudarski, Lisa Glazenburg, Eugene Berezikov

## Abstract

The evolutionary arms race between transposable elements (TEs) and their hosts contributes to genomic complexity. As TEs mobilization is deleterious for individual cells and organisms, their activity is restricted. During stress, TEs can be reactivated; however, the exact mechanisms vary. We discovered that in the flatworm *Macrostomum lignano*, LTR retrotransposons hijack the heat shock response pathway to boost their transcription at elevated temperatures. While it has been well-described in cruciferous plants, this is the first report of this mechanism in animal LTR retrotransposons. Our results suggest a convergent evolution of the heat stress response in LTR retrotransposons from animals and plants.

## Background

Eukaryotic genomes contain numerous families of diverse transposable elements (TEs) that are genetic units capable of self-replication and/or genomic location changes [1]. On a large scale, TEs movement and amplification have driven the evolution of genomes by expanding genetic diversity and creating complex regulatory landscapes and gene expression networks [1–5]. On the scale of an individual cell, organism, or even a small population, TEs activity is largely deleterious and, thus, is tightly controlled by internal factors in the host genome, such as DNA methylation and small RNA interference [6,7]. The silencing mechanisms must be relaxed during embryonic development or perturbed during the response to acute stressors such as infections, wounding, hybridization, changing temperature, and osmotic changes [6–13]. This lack of control creates opportunities for TEs expression and propagation; however, the exact activation mechanisms vary and remain poorly understood.

Previously, we computationally analyzed the diversity of Long Terminal Repeat (LTR) retrotransposons (LTR-RTs) in the regenerative flatworm model *Macrostomum lignano* [14,15]. Unexpectedly, we found canonical heat shock elements (HSE) in the promoters of some LTR-RT lineages [15]. HSEs are conserved binding sites for heat shock factor proteins that activate an evolutionary conserved cell protective mechanism known as heat shock response [16–18]. Based on the rigorously validated case of direct transcriptional activation of HSE-containing *ONSEN* LTR-RTs in *Arabidopsis thaliana* [19,20], we speculated that HSEs in *M. lignano* LTR-RTs independently evolved to gain transcriptional advantage compared to other genes during host defense machinery disruption after heat stress [15]. Here, we present the first experimental evidence of the direct transcriptional activation of flatworm LTR-RTs by heat stress. This represents one of the two known cases of this mechanism reported in animal TEs [21] and corroborates our hypothesis of interkingdom convergent evolution between plant and animal LTR-RTs.

## Results and Discussion

We revisited our previous LTR-RT annotation pipeline [15] to ensure more sensitive and specific detection of LTR-RTs with HSEs in *M. lignano*. For this, we utilized the newly developed DARTS algorithm [22] and combined it with stringent selection and clustering criteria to avoid over-grouping distantly related LTR-RT copies. Using this new pipeline, we identified 3077 LTR-RT copies, of which 1928 were full-length with intact terminal repeats. These were then sub-clustered into 455 LTR-RT lineages based on nucleotide sequence similarity. 12 lineages have one or more canonical HSE motifs in both of their LTRs (Additional file 1: Table S1).

To check whether the HSE-containing LTR-RTs in *M. lignano* transcriptionally respond to acute heat stress, we performed bulk RNA-seq experiments comparing worms placed for 2 h at either 37°C or 20°C. Upon heat stress, 2384 genes and 21 LTR-RT lineages (False Discovery Rate (FDR) < 0.05, log_2_FC > 1) were differentially expressed (Fig. 1A; Additional file 2: Table S2). A total of 1504 genes and seven LTR-RTs were downregulated, whereas 880 genes and 14 LTR-RTs were upregulated. Strikingly, we found that six of these LTR-RT lineages were strongly upregulated (log_2_FC = 7.7 ± 0.8, MEAN ± STDEV), matching or even exceeding the expression levels of the host’s heat shock protein genes (HSPs). Similar to HSPs, the six LTR-RT lineages contained one to four canonical HSE motifs upstream of transcription start sites (Fig. 1A-C). The other eight upregulated LTR-RTs showed a more moderate increase in expression (log_2_FC = 1.5 ± 0.6, MEAN ± STDEV) and did not contain HSEs (Fig. 1A, B; Additional file 2: Table S1, S2). The six LTR-RT lineages with HSEs that were not differentially expressed have only one HSE motif per LTR. This suggests the importance of motif location and/or number relative to the transcription start site sequence to induce transcriptional upregulation upon heat stress.

**Fig. 1.**
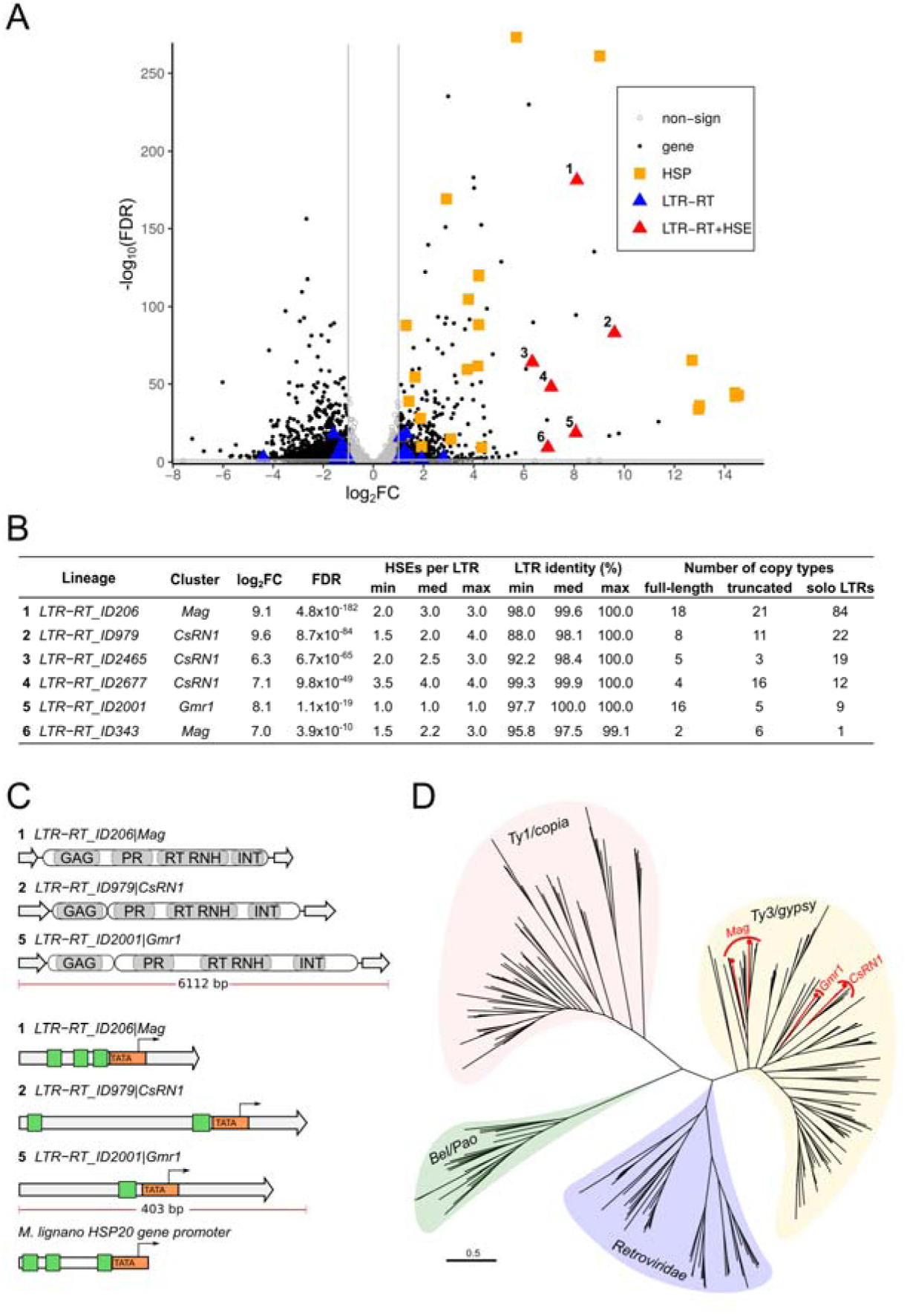
Transcriptional heat stress response and diversity of *M. lignano* HSE-containing LTR-RTs. **A** Volcano plot summarizing RNA-seq of transcriptional changes upon heat stress induction. Gray lines indicate the differential expression thresholds (FDR < 0.05, log2FC > 1). Non-sign: genes and LTR-RTs that were not differentially expressed. The numbers to the left of the red triangles of the LTR-RTs with HSEs correspond to those in Fig. 1B and C. **B** Detailed summary of differential expression, number of HSEs, and copy number of strongly upregulated LTR-RT lineages. Lineage names were defined based on their representative sequences. Clusters correspond to Fig. 1D. **C** Schematic structure of selected strongly upregulated HSE-containing LTR-RTs, their conserved protein domains, and their promoter regions. The general structures are shown at the top of the figure. Grey arrows: long terminal repeats (LTRs). LTRs schemes were compared to the *M. lignano Hsp20* promoter region (bottom panel). Oval-shaped blocks: open reading frames. GAG: nucleocapsid domain. PR: protease. RT: reverse transcriptase. RNH: ribonuclease H. INT: integrase. Orange arrow-shaped blocks: predicted TATA-boxes. Thin black arrows indicate transcription start sites. Green boxes: HSEs. (D) Schematic representation of the maximum-likelihood phylogenetic reconstruction of known LTR-RTs based on concatenated amino acid sequences of their RT, RH, and INT domains. Major families of LTR-RTs are highlighted in color, and their names are indicated inside. Red branches and cluster names indicate the positions of HSE-containing *M. lignano* LTR-RTs (Fig. 1A, B). Scale bar: average amino acid substitution rate per site. See Additional file 3: Fig. S2 for detailed phylogenetic analysis with clustering coefficients and LTR-RT names.

We confirmed the RNA-seq data by qRT-PCR and determined that as little as 5-15 min to 37°C exposure causes a strong transcriptional upregulation of two HSE-containing LTR-RTs (Additional file 3: Fig. S1). *M. lignano* dwells in the sand of intertidal zones in Mediterranean [14], and short heat exposures are likely to happen in this environment. Moreover, we previously showed that even milder temperatures of 30-35°C already caused a noticeable heat shock response in *M. lignano* [23]. Taken together, this suggests that on the evolutionary time scale, there could be a niche for LTR-RTs to harness *M. lignano’s* heat stress response machinery.

To date, the only rigorously validated case of LTR-RTs gaining direct heat sensitivity through the inclusion of HSEs upstream of the LTR promoters was ONSEN *Ty1/copia* LTR-RTs from cruciferous land plants (Brassicaceae), which includes the model species *Arabidopsis thaliana* [19,20,24]. In *M. lignano,* the six strongly upregulated HSE-containing LTR-RT lineages belong to a different major phylogenetic family of LTR-RTs, *Ty3/gypsy.* More specifically, they clustered within known animal-specific clades such as *Mag*, *CsRN1,* and *Gmr1* (Fig. 1D; Additional file 3: Fig. S2). The vast majority of *M. lignano* LTR-RTs were *Ty3/gypsy* from the *Mag* clade. Interestingly, only three out of 221 *Mag,* three out of 24 *CsRN1,* and one out of 115 *Gmr1* lineages in *M. lignano* had HSEs, indicating their later independent specialization (Additional file 1: Table S1). Therefore, *ONSEN*-like elements from Brassicaceae plants and HSE-containing LTR-RTs from *M. lignano* have convergently evolved HSE motifs in the LTRs.

In *A. thaliana*, the LTR sequence of *ONSEN* drives reporter transgene expression during heat stress [19]. This further proves that the transcriptional upregulation of *ONSEN* is not a result of the context of its copy insertion within the genome. Since transgenesis is available for *M. lignano* [25–27], we decided to adapt the *Arabidopsis* experiment to validate whether the isolated LTR promoter of the top upregulated *M. lignano* lineage (*Mlig_LTR-RT_ID206|Mag*) could drive mScarlet-I fluorescent protein expression *in vivo* (Fig. 2). We compared the performance of the generated NL52 *M. lignano* transgenic line during heat stress to the previously established and validated NL28 line, where the mScarlet-I coding gene is under the control of the *Mlig_Hsp20* promoter [23]. Both lines were derived from similarly organized transgenic constructs, where the elongation factor 1 alpha constitutive promoter drives mNeonGreen expression for the positive selection of transgenic animals (Fig. 2A). At 16 h post heat treatment (2 h at 37°C), red fluorescence was observed throughout the worms’ bodies in both transgenic lines, whereas it was undetectable in animals kept at 20°C (Fig. 2B). We did not observe transgene expression in the testes and ovaries of NL52 worms, in contrast to the strong signal in the ovaries of NL28 worms. This may indicate active silencing of the LTR-RT promoter in the germline or may be a result of the positional effect of transgene insertion due to random integration, which was previously observed for the ubiquitous *elongation factor 1 alpha* promoter in *M. lignano* [26]. We also investigated whether the *Mlig_LTR-RT_ID206|Mag* promoter could drive transgene expression during regeneration (Fig. 2C). *M. lignano* can restore all tissues and organs, including the gonads, posterior of its pharynx due to presence of a pluripotent population of somatic stem cells [28,29]. Theoretically, this could provide a post-embryonic window of opportunity where the germline can inherit random mutations and new TE insertions from somatic stem cells [9,30]. Interestingly, we observed an mScarlet-I signal in the blastema region of amputated animals (Fig. 2C); however, further research is required to confirm its expression in pluripotent stem cells and not only in lineage-restricted somatic progenitors.

**Fig. 2.**
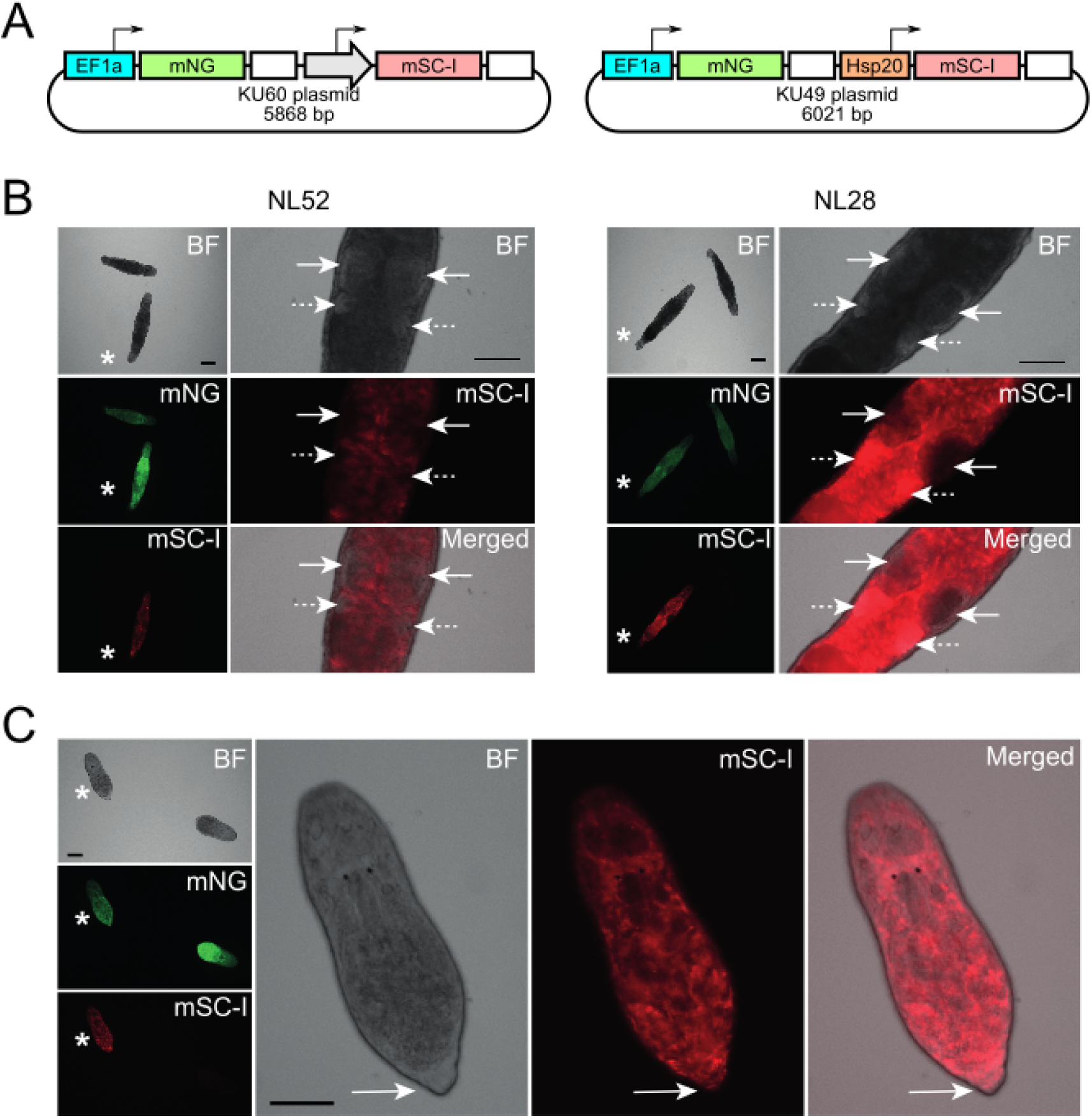
The isolated LTR sequence of *Mlig_LTR-RT_ID206|Mag* drives the transgenic reporter expression during heat stress. **A** Schematics of the donor plasmids KU60 and KU49 used to generate transgenic lines NL52 and NL28, respectively. Blocks with thin arrows above denote the promoters with 5’UTR regions, and the directions of the arrows reflect the orientation of a gene cassette in the plasmids. The 3’UTR regions of the elongation factor 1 alpha gene (EF1a) are shown as white blocks. The gray arrow-shaped block corresponds to the 5’ LTR of *Mlig_LTR-RT_ID206|Mag.* mNG: mNeonGreen, mSC-I: mScarlet-I. The full plasmid structure and sequences are shown in Additional file 3: Fig. S3. **B** Live imaging images of *M. lignano* transgenic worms before (not marked) and after heat shock (marked with asterisks). BF - brightfield. Scale bars: 100 μm. On the left side, the small panels show the comparisons. Close-up images of gonads of the corresponding heat-treated worms are shown on the right. Solid white arrows indicate the testes. Dashed white arrows indicate the ovaries. **C** Live images of regenerating *M. lignano* transgenic NL52 worms cut anterior to the testes before (not marked) and after heat shock (marked with asterisks). BF: brightfield. Scale bars: 100 μm. A close-up of the corresponding heat-treated worm is shown on the right side. Solid white arrows indicate the tail blastema region.

Numerous studies in eukaryotes have reported transcriptional activation of TEs due to heat stress [11,12,20,21,31]. However, in most cases, there is no data to explain the underlying molecular mechanisms explaining these observations. Loss of TEs’ silencing can be a result of indirect non-specific causes such as chromatin remodeling or co-transcription from promoters of neighboring genes [32,33]. Our discovery of HSE-containing LTR-RTs in the flatworm *M. lignano* is the second validated case of HSE acquisition by TEs in animals and the second report of this adaptation in LTR-RTs across the eukaryotic tree of life. The first case of HSE-derived heat sensitivity in animal TEs was shown in the rolling-circle DNA transposon *Helitron1_CE* from the nematode *Caenorhabditis elegans* [21,32]. HSEs in the promoter of *Helitron1_CE* provide a strong and direct Heat Shock Factor 1 dependent transcriptional upregulation of the transposon. Three independent events of the transcriptional heat activation mechanism across different TE families from divergent animal and plant taxa indicate that if searched, more TEs with HSEs can be discovered.

## Conclusions

We experimentally validated that heat stress strongly and directly upregulated the transcription of at least six HSE-containing *Ty3/gypsy* LTR-RTs in the flatworm *M. lignano*. This is the first example of HSEs’ acquisition by LTR-RTs in animals, and it is strikingly reminiscent of the *ONSEN Ty1/copia* lineage from Brassicaceae plants. The recurrent emergence of the HSEs direct heat activation mechanism in divergent families of LTR-RTs and their hosts suggests that there may be more such events across the tree of life.

## Methods

### M. lignano lines and culturing

Culturing of *Macrostomum lignano* was done as previously described [34]. Wild type NL12 worms were kept at 20°C. The transgenic lines NL28 and NL52 were initially kept at 25°C for faster propagation and then moved to 20°C for maintenance of the cultures. The NL28 line was generated in a previous study [23]. The NL52 line was made in this study. A detailed definition of the expressed transgenes can be found in the Results and Discussion section and Additional file 3: Fig. S3.

### Computational mining, annotation, and clustering of LTR-RTs

We used the genome assembly Mlig_3_7 (GenBank accession number: GCA_002269645.1) [25]. DARTS with default parameters was used to mine LTR-RTs, annotate their domain composition, and identify LTRs [22]. To cluster nucleotide sequences of LTR-RTs into lineages, we used MMseqs2 [35] with the following command line parameters: “easy-cluster --min-seq-id 0.7 --min-aln-len 80 --cov-mode 0 -c 0.4 --cluster-mode 1”. Specifically, we required 70% minimum sequence identity, with 40% bidirectional coverage. Cluster representatives were selected based on maximum DARTS completeness score [22]. LTR-RT copies were considered complete if they had predicted LTRs and GAG, PR, RT, RH, and INT protein domains. Truncated or damaged copies lack at least one of these features. To better estimate the number of truncated copies and solo-LTRs (inter-copy homologous recombination by-products) for selected lineages, we re-mapped the cluster representative sequences to the genome using BLAST [36]. Regions were counted based on their alignment either to only LTRs, the internal portion between LTRs, or the full LTR-RT sequence. Preliminary classification of the lineages was done based on 1-to-1 nucleotide BLAST identity to previously phylogenetically classified LTR-RT copies [15].

### HSEs and TATA-box annotation, phylogenetic analysis

Identification of HSEs and phylogenetic analyses of the HSE-containing lineages were done as previously described [15]. Putative TATA-box sequences were predicted using the Neural Network Promoter Prediction tool available at https://www.fruitfly.org/seq_tools/nnppHelp.html.

### RNA-seq library preparation and differential expression analysis

NL12 worms were starved for 48 h, changing the sea water medium after first 24 h and right before the experiment. Total RNA was extracted using TRIzol™ Reagent (Invitrogen) from eight batches of 100 worms kept either at 20°C (four replicates) or at 37°C (four replicates) for 2 h. Right before the extraction, worms were put on ice for 5 min to assist the collection of animals and the sea water medium. 3’ end RNA-seq libraries were prepared using the Smart-3SEQ protocol from 100 ng of total RNA input as described in the original publication [37]. The libraries were pooled together and sequenced on an Illumina NextSeq 500 instrument at the ERIBA Research Sequencing Facility (Groningen, the Netherlands). Raw reads were trimmed with TrimGalore v.0.6.7 [38] and deduplicated with SeqKit v.2.4.049 [39]. Then they were mapped to the *M. lignano* genome assembly Mlig_3_7 using STAR 2.7.9a [40]. Gene counts were generated with the TEToolkit 2.0.3 [41] using the latest published *M. lignano* transcriptome assembly and annotation Mlig_RNA_3_7_DV1.v3 [25,42] and the GTF annotation file of LTR-RT representatives’ copies insertions was generated by DARTS. Subsequent differential gene expression analysis was performed with the DEseq2 package in R [43].

### qRT-PCR

Wild type NL12 worms were put at 37°C for 0, 5, 15, 30, 60, and 90 min. The conditions were repeated 3 times, using 100 worms per replicate. RNA extraction was performed as described above, but total RNA samples were additionally treated with the DNA-free™ DNA Removal Kit (Invitrogen) to remove gDNA traces. Reverse transcription and Quantitative RT-PCR was done using the Light Cycler 480 (Roche) as previously described [28] with primers specific for *Mlig_LTR-RT_ID206|Mag, Mlig_LTR-RT_ID2465|CsRN1,* and *Mlig_Hsp20* as well as *Mlig_Gapdh* as a reference target gene (Additional file 4: Table S3).

### Transgenic constructs and transgenesis

The NL28 line was generated previously using random integration transgenesis through KU#49 plasmid microinjections into single cell stage *M. lignano* eggs [23,25]. The plasmid KU#60 was generated from KU#49 by swapping the *Mlig_Hsp20* promoter with the LTR promoter of *Mlig_LTR-RT_ID206|Mag* (Additional file 3: Fig. S3). The LTR sequence was PCR amplified from *M. lignano* genomic DNA using primers specified in Additional file 4: Table S3. Microinjections and selection of transgenic animals was done as previously described [25,26].

### Microscopy and imaging

Live imaging of *M. lignano* transgenic lines was performed using a Zeiss Axio Zoom V16 epifluorescent stereomicroscope with an HRm digital camera and Zeiss filter sets 38HE (FITC) and 43HE (DsRed). The worms were prepared as described before [26,34]. Two animals, control and heat-treated, were put and imaged together in one field of view. The exposure settings were kept the same for the NL52 and NL28 transgenic lines.

## Supplementary Information

**Additional file 1: Table S1.** Summary of LTR-RTs identified in the study.

**Additional file 1: Table S2.** Differentially expressed genes and LTR-RTs upon heat shock in *M. lignano*.

**Additional file 1: Fig. S1.** qRT-PCR analysis of sensitivity of two upregulated LTR-RT lineages transcription to time of heat stress. **Fig. S2.** LTR-RTs upregulated upon heat stress belong to the *Ty3/gypsy Mag*, *CsRN1*, and *Gmr1* clades. **Fig. S3.** KU60 plasmid scheme and its GenBank formatted annotation.

**Additional file 4: Table S3.** qRT-PCR and cloning primers.

## Declarations

### Ethics approval and consent to participate

Not applicable

## Consent for publication

Not applicable

## Availability of data and materials

All data generated or analyzed during this study are included in this published article and its supplementary information files. All plasmids and *M. lignano* worm lines generated in this study are available from the corresponding authors on request.

## Authors’ contributions

Conception of the study: K.U. and E.B.; data analysis: K.U., M.B., and E.B.; transgenesis: K.U. and J.W.; microscopy: S.M.; animal culture and RNA-seq library preparation: K.U., S.M., and L.G.; manuscript writing: K.U., S.M., and E.B.; funding acquisition: E.B. All authors have read and approved the manuscript.

