## Additional file 3: Fig. S for "Heat stress drives transcription of LTR retrotransposons in the regenerative flatworm *Macrostomum lignano*"

#### Supplementary Figure Legends

**Fig. S1** qRT-PCR analysis of sensitivity of the top two upregulated LTR-RT lineages transcription to time of heat stress. LTR-RT names correspond to those in Fig. 1. Expression values are relative to the 0 min time point. *Mlig_Hsp20* gene was used as a positive control. Expression values between samples were normalized to GAPDH.

**Fig. S2** LTR-RTs upregulated upon heat stress belong to the *Ty3/gypsy Mag*, *CsRN1,* and *Gmr1* clades. Unrooted maximum likelihood tree of phylogenetic relationships among representatives of four major LTR-RTs groups reconstructed based on concatenated amino-acid sequences of their RT, RNH and INT protein domains. Branch support values (aLRT/UFBoot) are indicated above the branches leading to corresponding cluster. Major phylogenetic groups are highlighted with coloured blocks. Borders of phylogenetic clusters are drawn as vertical black bars to the right from the tree. Names of the clusters and LTR-RTs are according to GyDB (http://gydb.org). Names of the *M. lignano* HSE-containing LTR-RT lineage representatives and the clusters they belong to are highlighted in red.

**Fig. S3** KU60 plasmid scheme and its GenBank formatted annotation. An annotation of the LTR sequence with Heat Shock Elements of the *Mlig_LTR-RT_ID206|Mag* is provided before the GenBank annotation. The LTR promoter sequence of the *Mlig_LTR-RT_Mag|ID206 LTR-RT*. HSE motifs are in bold and highlighted in yellow. Putative TATA-box is highlighted in blue.

Supplementary Figure 1

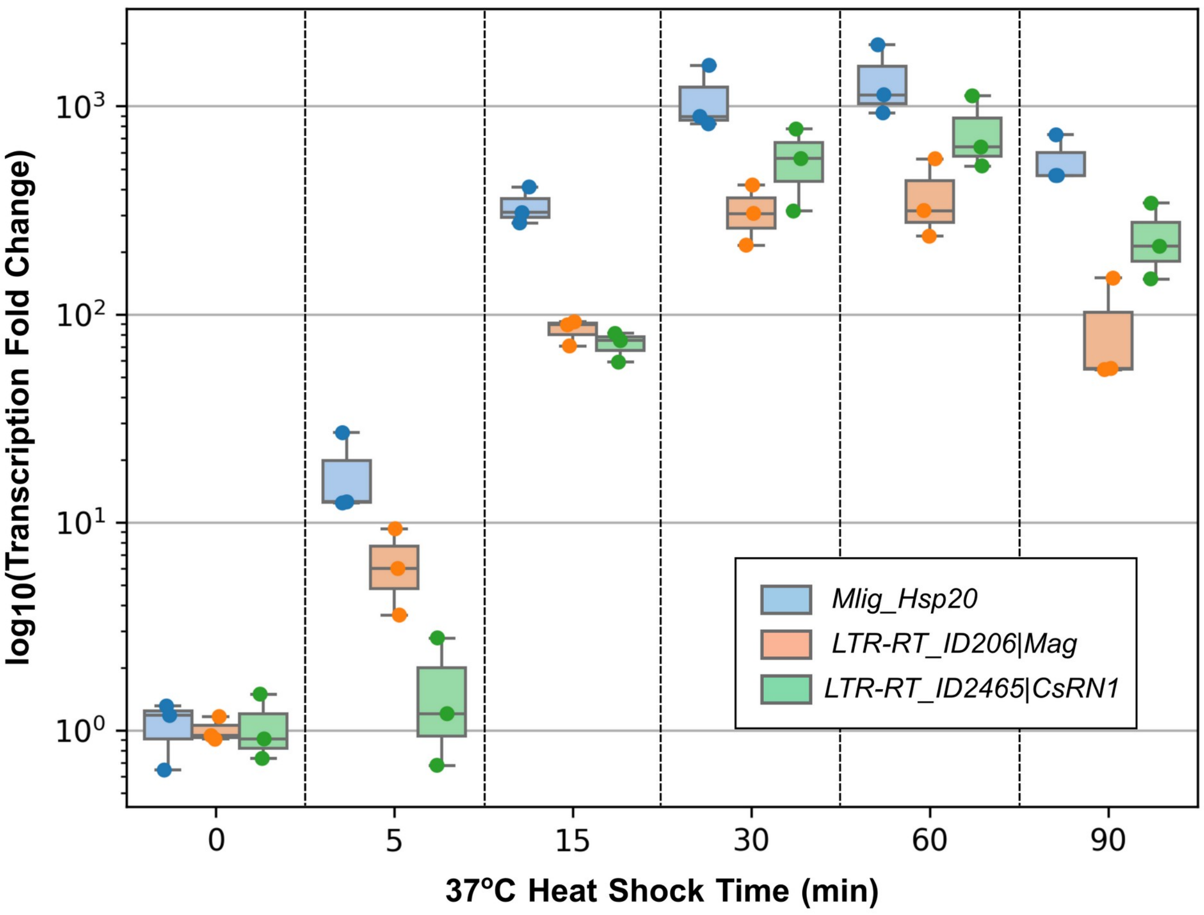

Supplementart Figure 2

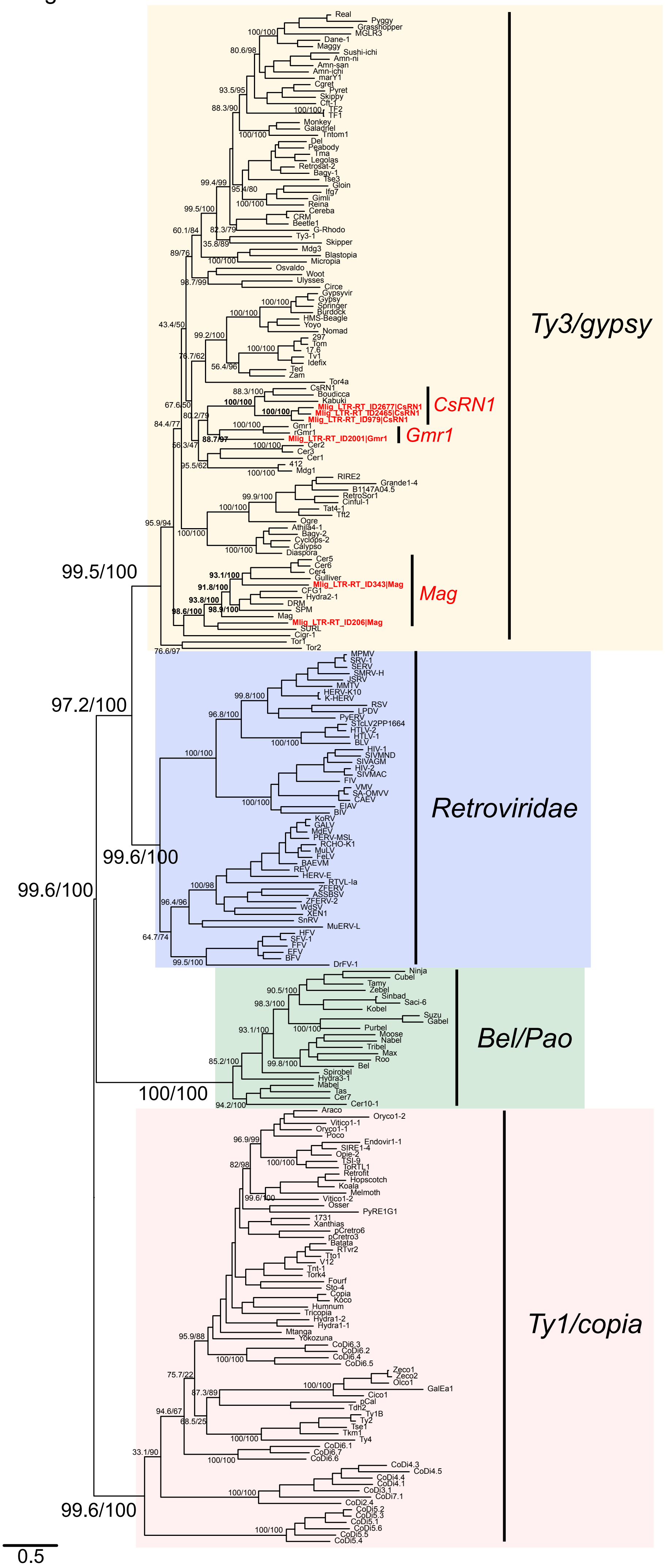

Supplementary Figure 3

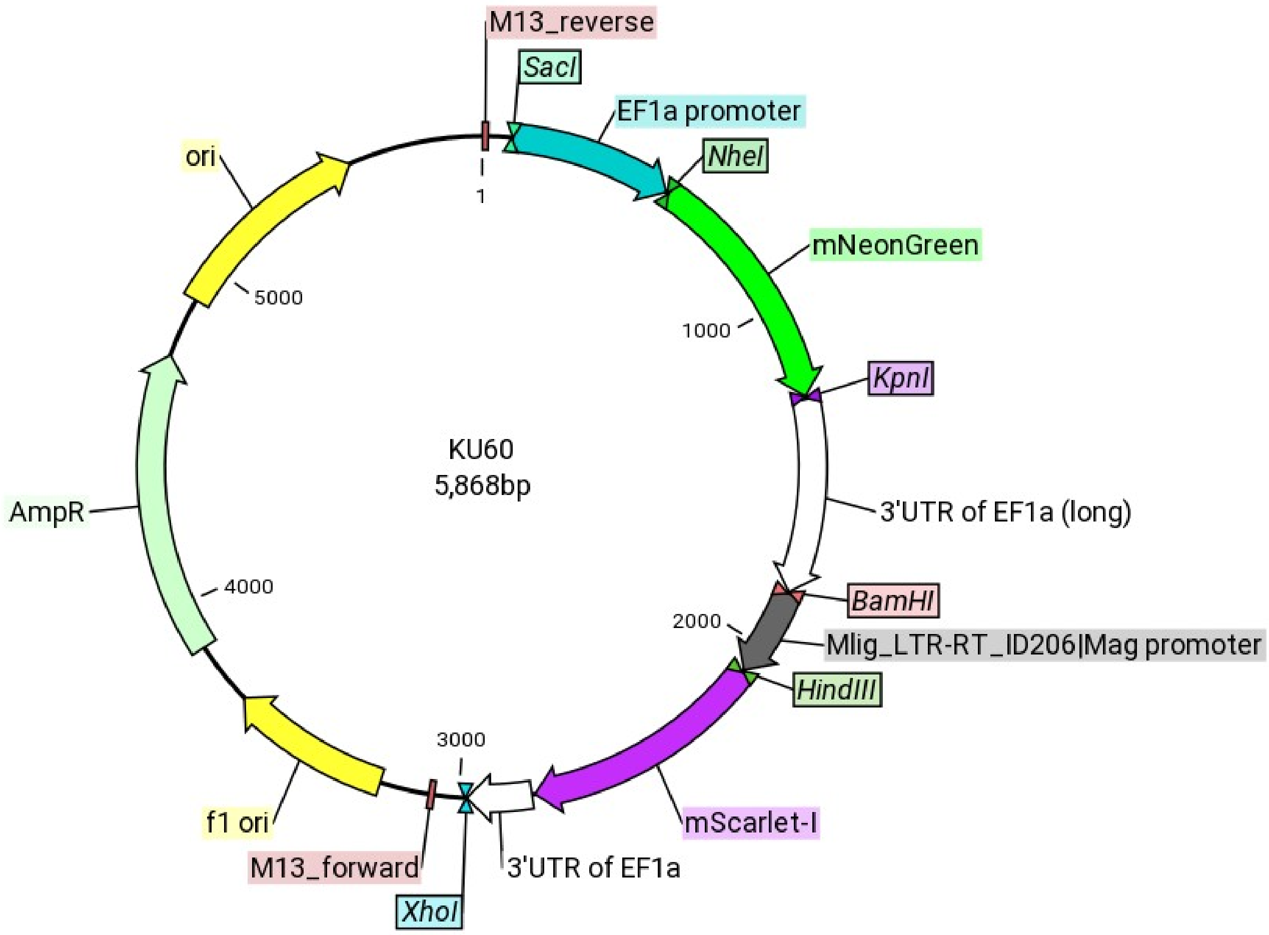


**5′** [BamHI]
TGTTGTGTATTGCAAAGGCATGCGATGCGTAGGGGTAGTTCGGGGACAGCGT**TTC**TA**GAA**AC**TTC**CACCGCGGACTTGTT
TACGCATAG**TTC**CA**GAA**GT**TTC**CTTCCCCGGTCT**TTC**TA**GAA**CT**TTC**CTTCCCCGCGTGCATAAATATACGAGCCGCCCT
CCTATTGGCCAAGCCCCTCTTTTGGTGCCACTGAGCGCCCGGATAATTATCTATCTTGTTTCTGCGTTCGTTAATTCAAC
AACAGAACAATG[HindIII] **3′**

### GENE BANK FORMATTED PLASMID ANNOTATION

```
LOCUS       KU60       5868 bp    DNA     circular UNA 17-Dec-2024
DEFINITION
FEATURES                 Location/Qualifiers
     primer              2..18
                         /label="M13_reverse"
     EF1a                90..551
                         /label="EF1a promoter"
     mNeonGreen          561..1265
                         /label="mNeonGreen"
     3'UTR               1275..1827
                         /label="3'UTR of EF1a (long)"
     LTR_promoter        1834..2082
                         /label="Mlig_LTR-RT_ID206|Mag promoter"
     mScarlet-I          2092..2784
                         /label="mScarlet-I"
     3'UTR               2794..2977
                         /label="3'UTR of EF1a"
     primer              complement(3070..3086)
```

```
                     /label="M13_forward"
     ori             3227..3682
                     /label="f1 ori"
     AmpR            3865..4725
                     /label="AmpR"
     ori             4896..5484
                     /label="ori"
ORIGIN
        1 ACAGGAAACA GCTATGACCA TGATTACGCC AAGCTATTTA GGTGACACTA TAGAATACTC
       61 AAGCTATGCA TCCAACGCGT TGGGAGCTCC GTCTTCTGTT CAGTGTTACA AGTTTTGATG
      121 TAAAAAAACAA TATTGGCCTA TAACATTTTC ATATATCCTC TCTACATTTT CATATATTTT
      181 AAAAAGTATT TAAAAAACAT CTTAAAAGTA TTCCATTTTA TTATTAACTG AAAAACTAAC
      241 GTAAAATTAC AATATATCTC AAAACTTTCT AAAGTCGTAT ATTTTCCGGC ATCGTTAATT
      301 TTAACGACAA TCGTGAAAGT TCGTGAACAA GTTTGCTGTT CCTCACCTAA ATTTGAATTA
      361 CTATCGCTCT TGTCTGAATG GATTTCTTAC TGCAGATATC CTGAAGTAAG TCTTTCAATT
      421 TGTGAATTGT AAGTAAAGTA TTTTTTATTA CTGCTATCAT ATTTGTTGAA TTTGTTCCAA
      481 ATTACTATTT CTTGCTTTTA ATTCAGTTCC TTTTATTTAT TCCTAAACTC AGGTAGTTTA
      541 ACAACAGCAT TATGGCTAGC GTGAGCAAGG GCGAGGAGGA CAACATGGCC AGCCTGCCGG
      601 CCACCCACGA GCTGCACATC TTCGGCAGCA TCAACGGCGT GGACTTCGAC ATGGTGGGCC
      661 AGGGCACCGG CAACCCGAAC GACGGCTACG AGGAGCTGAA CCTGAAGAGC ACCAAGGGCG
      721 ACCTTCAGTT CAGCCCGTGG ATTCTGGTGC CGCACATCGG CTACGGCTTC CACCAGTACC
      781 TGCCGTACCC GGACGGCATG AGCCCGTTCC AGGCCGCTAT GGTGGACGGC AGCGGCTACC
      841 AGGTGCACCG CACCATGCAG TTCGAGGACG GCGCCAGCCT GACCGTGAAC TACCGCTACA
      901 CCTACGAGGG CAGCCACATC AAGGGCGAGG CCCAGGTGAA GGGCACCGGC TTCCCGGCCG
      961 ACGGCCCGGT GATGACCAAC AGCCTGACCG CCGCCGACTG GTGCCGCAGC AAGAAGACCT
     1021 ACCCGAACGA CAAGACCATC ATCAGCACCT TCAAGTGGAG CTACACCACC GGCAACGGCA
     1081 AGCGCTACCG CAGCACCGCC CGCACCACCT ACACCTTCGC CAAGCCGATG GCCGCCAACT
     1141 ACCTGAAGAA CCAGCCGATG TACGTGTTCC GCAAGACCGA GCTGAAGCAC AGCAAGACCG
     1201 AGCTGAACTT CAAGGAGTGG CAGAAGGCCT TCACCGACGT GATGGGCATG GACGAGCTGT
     1261 ACAAGGGTAC CTGAAATGAC CCGCACCTGT GTAATTCTGG AGGCATTGCA GTTAACTGAA
     1321 CTATTTGGAA TGCTCGATTG TCGATCAATA AAGAATGCGA AGTTAAAGTT GCGAGACTTG
     1381 TTTGTTTGGA AATTCCTGTC TCGATTTGTT TAATTTCTTT GTGTTTCAAT CTATTTGTTG
     1441 CTAGTATTTC CTGACTAAAG AGCGACGTAA AGCCCGAACT TCACAAATGT ATGGGCACAA
     1501 GTATAGGCTG CATAATGAAG TAGCCAGTTA ATTGCAACGA ATTAACAGTT CATTTATTCA
     1561 TATTACAAGG GGCGGGAACT GAAAAGTTGG GAGGCTGTAC TTTTTAATAA CATTGCTTCA
     1621 AGCATTACTT GAAGCAGTAT TTACCACAGA CTAAGTGACT GCAAAGGAAC ACTAACTCGT
     1681 TCTGTAAGTC CCTAAATTTG ATGTCTTCTG TTCAATTTCA AACAATAACT TGGAATCTCT
     1741 GGGTAAAATT CTGAATCATA CAATTGTCAG TATCAGTACA TGCTAACAAC TGAGTGTACA
     1801 TGCCAACTGT GGGCTAGTAT CAGCGAAGGA TCCTGTTGTG TATTGCAAAG GCATGCGATG
     1861 CGTAGGGGTA GTTCGGGGAC AGCGTTTCTA GAAACTTCCA CCGCGGACTT GTTTACGCAT
     1921 AGTTCCAGAA GTTTCCTTCC CCGGTCTTTC TAGAACTTTC CTTCCCCGCG TGCATAAATA
     1981 TACGAGCCGC CCTCCTATTG GCCAAGCCCC TCTTTTGGTG CCACTGAGCG CCCGGATAAT
     2041 TATCTATCTT GTTTCTGCGT TCGTTAATTC AACAACAGAA CAATGAAGCT TGTGAGCAAG
     2101 GGCGAGGCCG TGATCAAGGA GTTCATGCGC TTCAAGGTGC ACATGGAGGG CAGCATGAAC
     2161 GGCCACGAGT TCGAGATCGA GGGCGAGGGC GAGGGCCGCC CGTACGAGGG CACCCAGACC
     2221 GCCAAGCTGA AGGTGACCAA GGGCGGCCCG CTGCCGTTCA GCTGGGACAT CCTGAGCCCG
     2281 CAGTTCATGT ACGGCAGCCG CGCCTTCATC AAGCACCCGG CCGACATCCC GGACTACTAC
     2341 AAGCAGAGCT TCCCGGAGGG CTTCAAGTGG GAGCGCGTGA TGAACTTCGA GGACGGCGGC
     2401 GCCGTGACCG TGACCCAGGA CACCAGCCTG GAGGACGGCA CCCTGATCTA CAAGGTGAAG
     2461 CTGCGCGGCA CCAACTTCCC GCCGGACGGC CCGGTGATGC AGAAGAAGAC TATGGGCTGG
     2521 GAGGCCAGCA CCGAGCGCCT GTACCCGGAG GACGGCGTGC TGAAGGGCGA CATCAAGATG
     2581 GCCCTGCGCC TGAAGGACGG CGGTCGCTAC CTGGCCGACT TCAAGACCAC CTACAAGGCC
     2641 AAGAAGCCGG TGCAGATGCC GGGCGCCTAC AACGTGGACC GCAAGCTGGA CATCACCAGC
     2701 CACAACGAGG ACTACACCGT GGTGGAGCAG TACGAGCGCA GCGAGGGCCG CCACAGCACC
     2761 GGCGGCATGG ACGAGCTGTA CAAGTCTAGA TGAGCAATGA CCCGCACCTG TGTAATTCTG
     2821 GAGGCATTGC AGTTAACTGA ACTATTTGGA ATGCTCGATT GTCGATCAAT AAAGAATGCG
     2881 AAGTTAAAGT TGCGAGACTT GTTTGTTTGG AAATTCCTGT CTCGATTTGT TTAATTTCTT
     2941 TGTGTTTCAA TCTATTTGTT GCTAGTATTT CCTGACTCTC GAGATAATCG AATTCCCGCG
     3001 GCCGCCATGG CGGCCGGGAG CATGCGACGT CGGGCCCAAT TCGCCCTATA GTGAGTCGTA
     3061 TTACAATTCA CTGGCCGTCG TTTTACAACG TCGTGACTGG GAAAACCCTG GCGTTACCCA
     3121 ACTTAATCGC CTTGCAGCAC ATCCCCCTTT CGCCAGCTGG CGTAATAGCG AAGAGGCCCG
     3181 CACCGATCGC CCTTCCCAAC AGTTGCGCAG CCTGAATGGC GAATGGACGC GCCCTGTAGC
     3241 GGCGCATTAA GCGCGGCGGG TGTGGTGGTT ACGCGCAGCG TGACCGCTAC ACTTGCCAGC
     3301 GCCCTAGCGC CCGCTCCTTT CGCTTTCTTC CCTTCCTTTC TCGCCACGTT CGCCGGCTTT
```

```
3361 CCCCGTCAAG CTCTAAATCG GGGGCTCCCT TTAGGGTTCC GATTTAGTGC TTTACGGCAC
3421 CTCGACCCCA AAAAACTTGA TTAGGGTGAT GGTTCACGTA GTGGGCCATC GCCCTGATAG
3481 ACGGTTTTTC GCCCTTTGAC GTTGGAGTCC ACGTTCTTTA ATAGTGGACT CTTGTTCCAA
3541 ACTGGAACAA CACTCAACCC TATCTCGGTC TATTCTTTTG ATTTATAAGG GATTTTGCCG
3601 ATTTCGGCCT ATTGGTTAAA AAATGAGCTG ATTTAACAAA AATTTAACGC GAATTTTAAC
3661 AAAATATTAA CGCTTACAAT TTCCTGATGC GGTATTTTCT CCTTACGCAT CTGTGCGGTA
3721 TTTCACACCG CATCAGGTGG CACTTTTCGG GGAAATGTGC GCGGAACCCC TATTTGTTTA
3781 TTTTTCTAAA TACATTCAAA TATGTATCCG CTCATGAGAC AATAACCCTG ATAAATGCTT
3841 CAATAATATT GAAAAAGGAA GAGTATGAGT ATTCAACATT TCCGTGTCGC CCTTATTCCC
3901 TTTTTTGCGG CATTTTGCCT TCCTGTTTTT GCTCACCCAG AAACGCTGGT GAAAGTAAAA
3961 GATGCTGAAG ATCAGTTGGG TGCACGAGTG GGTTACATCG AACTGGATCT CAACAGCGGT
4021 AAGATCCTTG AGAGTTTTCG CCCCGAAGAA CGTTTTCCAA TGATGAGCAC TTTTAAAGTT
4081 CTGCTATGTG GCGCGGTATT ATCCCGTATT GACGCCGGGC AAGAGCAACT CGGTCGCCGC
4141 ATACACTATT CTCAGAATGA CTTGGTTGAG TACTCACCAG TCACAGAAAA GCATCTTACG
4201 GATGGCATGA CAGTAAGAGA ATTATGCAGT GCTGCCATAA CCATGAGTGA TAACACTGCG
4261 GCCAACTTAC TTCTGACAAC GATCGGAGGA CCGAAGGAGC TAACCGCTTT TTTGCACAAC
4321 ATGGGGGATC ATGTAACTCG CCTTGATCGT TGGGAACCGG AGCTGAATGA AGCCATACCA
4381 AACGACGAGC GTGACACCAC GATGCCTGTA GCAATGGCAA CAACGTTGCG CAAACTATTA
4441 ACTGGCGAAC TACTTACTCT AGCTTCCCGG CAACAATTAA TAGACTGGAT GGAGGCGGAT
4501 AAAGTTGCAG GACCACTTCT GCGCTCGGCC CTTCCGGCTG GCTGGTTTAT TGCTGATAAA
4561 TCTGGAGCCG GTGAGCGTGG GTCTCGCGGT ATCATTGCAG CACTGGGGCC AGATGGTAAG
4621 CCCTCCCGTA TCGTAGTTAT CTACACGACG GGGAGTCAGG CAACTATGGA TGAACGAAAT
4681 AGACAGATCG CTGAGATAGG TGCCTCACTG ATTAAGCATT GGTAACTGTC AGACCAAGTT
4741 TACTCATATA TACTTTAGAT TGATTTAAAA CTTCATTTTT AATTTAAAAG GATCTAGGTG
4801 AAGATCCTTT TTGATAATCT CATGACCAAA ATCCCTTAAC GTGAGTTTTC GTTCCACTGA
4861 GCGTCAGACC CCGTAGAAAA GATCAAAGGA TCTTCTTGAG ATCCTTTTTT TCTGCGCGTA
4921 ATCTGCTGCT TGCAAACAAA AAAACCACCG CTACCAGCGG TGGTTTGTTT GCCGGATCAA
4981 GAGCTACCAA CTCTTTTTCC GAAGGTAACT GGCTTCAGCA GAGCGCAGAT ACCAAATACT
5041 GTTCTTCTAG TGTAGCCGTA GTTAGGCCAC CACTTCAAGA ACTCTGTAGC ACCGCCTACA
5101 TACCTCGCTC TGCTAATCCT GTTACCAGTG GCTGCTGCCA GTGGCGATAA GTCGTGTCTT
5161 ACCGGGTTGG ACTCAAGACG ATAGTTACCG GATAAGGCGC AGCGGTCGGG CTGAACGGGG
5221 GGTTCGTGCA CACAGCCCAG CTTGGAGCGA ACGACCTACA CCGAACTGAG ATACCTACAG
5281 CGTGAGCTAT GAGAAAGCGC CACGCTTCCC GAAGGGAGAA AGGCGGACAG GTATCCGGTA
5341 AGCGGCAGGG TCGGAACAGG AGAGCGCACG AGGGAGCTTC CAGGGGGAAA CGCCTGGTAT
5401 CTTTATAGTC CTGTCGGGTT TCGCCACCTC TGACTTGAGC GTCGATTTTT GTGATGCTCG
5461 TCAGGGGGGC GGAGCCTATG GAAAAACGCC AGCAACGCGG CCTTTTTACG GTTCCTGGCC
5521 TTTTGCTGGC CTTTTGCTCA CATGTTCTTT CCTGCGTTAT CCCCTGATTC TGTGGATAAC
5581 CGTATTACCG CCTTTGAGTG AGCTGATACC GCTCGCCGCA GCCGAACGAC CGAGCGCAGC
5641 GAGTCAGTGA GCGAGGAAGC GGAAGAGCGC CCAATACGCA AACCGCCTCT CCCCGCGCGT
5701 TGGCCGATTC ATTAATGCAG CTGGCACGAC AGGTTTCCCG ACTGGAAAGC GGGCAGTGAG
5761 CGCAACGCAA TTAATGTGAG TTAGCTCACT CATTAGGCAC CCCAGGCTTT ACACTTTATG
5821 CTTCCGGCTC GTATGTTGTG TGGAATTGTG AGCGGATAAC AATTTCAC
//
```
